# Direct Speech Reconstruction from Sensorimotor Brain Activity with Optimized Deep Learning Models

**DOI:** 10.1101/2022.08.02.502503

**Authors:** Julia Berezutskaya, Zachary V. Freudenburg, Mariska J. Vansteensel, Erik J. Aarnoutse, Nick F. Ramsey, Marcel A.J. van Gerven

**Author notes:** these authors contributed equally to this work (Zachary V. Freudenburg, Mariska J. Vansteensel).

## Abstract

Development of brain-computer interface (BCI) technology is key for enabling communication in individuals who have lost the faculty of speech due to severe motor paralysis. A BCI control strategy that is gaining attention employs speech decoding from neural data. Recent studies have shown that a combination of direct neural recordings and advanced computational models can provide promising results. Understanding which decoding strategies deliver best and directly applicable results is crucial for advancing the field. In this paper, we optimized and validated a decoding approach based on speech reconstruction directly from high-density electrocorticography recordings from sensorimotor cortex during a speech production task. We show that 1) dedicated machine learning optimization of reconstruction models is key for achieving the best reconstruction performance; 2) individual word decoding in reconstructed speech achieves 92-100% accuracy (chance level is 8%); 3) direct reconstruction from sensorimotor brain activity produces intelligible speech. These results underline the need for model optimization in achieving best speech decoding results and highlight the potential that reconstruction-based speech decoding from sensorimotor cortex can offer for development of next-generation BCI technology for communication.

## Introduction

Due to a motor neuron disease or a brainstem stroke, people can lose all voluntary control over their muscles, including the ability to speak or use other movements for communication. Brain-computer interface (BCI) technology aims to provide a means of communication to these individuals. Recent advances in BCI research have demonstrated the potential of current approaches to record, process and analyse neural activity in order to decode speech^1–28^. This work is primarily conducted in able-bodied human subjects who participate in speech production experiments while their brain activity is recorded, with the goal to infer spoken speech from the acquired brain signals. This setup is used as a testbed for development and validation of speech decoding frameworks prior to their use in real-world BCI applications with paralyzed individuals^29–34^. See^35–45^ for reviews on BCIs for communication and speech.

Previous work demonstrates that speech decoding from brain activity is a challenging task. In order to realize a real-world speech-based BCI application, several key components of such a system need to be identified. This includes decisions on the neural recording modality (acquisition technique), the cortical areas that are most informative for decoding (neural sources), the target speech features to decode (decoding targets) and the decoding model itself. Regarding the acquisition technique, the most promising results in the field so far appear to have come from the use of invasive recording modalities, such as electrocorticography (ECoG)^10–12, 15, 18, 20, 23, 24^. This is because ECoG grids, and particularly those with a high spatial density (high-density ECoG; HD ECoG), provide excellent signal quality and high spatial and temporal resolution, which are key for achieving accurate decoding results. Preparatory BCI research and development can be performed in able-bodied patients with medication-resistant epilepsy who undergo temporarily implantation with ECoG electrodes (typically, for 7-10 days) for clinical monitoring of their condition and potential subsequent removal of neural sources of epilepsy. Regarding the neural sources, among the areas most informative for decoding is the motor cortex, which coordinates voluntary muscle control. Studies in amputees and severely paralyzed individuals have shown that motor cortex activity during attempted movement of the missing limb or attempted movement by the paralyzed individual is similar to the activity during actual movement in able-bodied individuals^31, 46^. Motor-based BCI research has led to the development of various BCI applications, such as the control of a robotic arm in tetraplegia^47^, a BCI speller based on attempted hand movement in a locked-in individual^31^ and a BCI for speech detection and decoding in an individual with anarthria^33^.

Choices of decoding targets and decoding model, however, are more difficult and varied. In principle, use of motor cortex as the neural source for decoding means that one of the most straightforward targets of decoding would be information about the movement of facial muscles involved in speech production. This is typically referred to as the kinematic articulator traces^18, 48^. However, for development of such speech decoding approach, articulator movements need to be recorded during speech simultaneously with the brain activity, which is rarely feasible in ECoG research due to the additional burden it would inflict on the study participants. Moreover, another system will be required to map the decoded kinematic movements to language information, such as phonemes, syllables or words^49^. Instead, one could consider using these language labels as targets of decoding. However, such labels are slow-changing speech features that have not been trivial to decode accurately from the brain activity in the past, especially when decoding a large number of classes^10, 30^. An alternative candidate for the target of speech decoding is acoustic speech features from simultaneously acquired microphone recordings of produced speech, available at the BCI development stage that relies on experiments in able-bodied participants^20, 21, 26, 27^. Such acoustic properties are fast-changing, highly repeatable speech features that have been previously shown to explain patterns of motor and premotor brain activity^50, 51^. Importantly for BCI, learning to infer just a handful of such features from brain signals could open up the possibility of open-vocabulary speech reconstruction. While previous work has shown that acoustic information is a promising candidate for decoding^20, 21, 26, 27^, it remains unclear to what extent acoustic properties of speech may be decoded directly from sensorimotor cortex using HD ECoG recordings and how intelligible resulting reconstructed speech would be.

A number of speech decoding models from brain activity have been proposed in the past. Recently, deep learning models have become increasingly more popular in the BCI field due to their potential for learning complex relationships between sources and targets of decoding^18, 20, 24–26, 33, 52^. However, there is a plethora of deep learning architectures^53^ and parameter choices. As a result, many studies tend to borrow model architectures and parameters from other fields and apply them with minimal changes to neural data for speech decoding. Thus far, no comprehensive study on optimization of deep learning models for speech reconstruction has been performed. Moreover, there is a lack of consensus regarding choices of brain and audio speech features that are used in such models. Current studies use different methods for ECoG data preprocessing, including de-noising reference schemes and frequency component ranges, as well as different types of audio speech features. Finally, evaluation of speech decoding models is also approached differently, and consensus regarding which metrics may be more informative is currently lacking. Overall, it remains unclear whether such a neuroengineering approach for speech reconstruction from the motor cortex could provide the basis for a real-world autonomous BCI application for decoding intended speech, and what the next practical steps for it would be.

In the present work, we aimed to decode acoustic properties of speech from sensorimotor cortex by building and evaluating deep learning models for the reconstruction of speech acoustics. For this, we collected HD ECoG data during a word production task, in which five human subjects spoke 12 unique words out loud, repeating each word ten times (Figure 1). We performed a dedicated machine learning procedure to optimize parameters of the speech reconstruction models, and examined metrics for evaluation of the reconstruction results (Figure 2). First, we report a significant improvement of the reconstruction accuracy due to model optimization and identify parameters associated with best results. Second, we show that more complex decoding models lead to better speech reconstruction. Third, we found that individual metrics for evaluation of speech reconstruction reflect different aspects of model performance and that low- and high-level metrics can be dissociated. Fourth, decoding of individual words from speech reconstructed from sensorimotor brain activity achieved 92-100% accuracy (chance level is 8%). Fifth, reconstructed speech exhibited high perceptual quality due to the model optimization procedure. Altogether, these results demonstrate the potential for reconstruction of identifiable, high-quality speech directly from sensorimotor brain activity and the importance of model optimization in achieving best perceptual results. These findings contribute to the state of the art in the field of speech decoding and reconstruction for BCI and have the potential to guide its further development.

**Figure 1.**
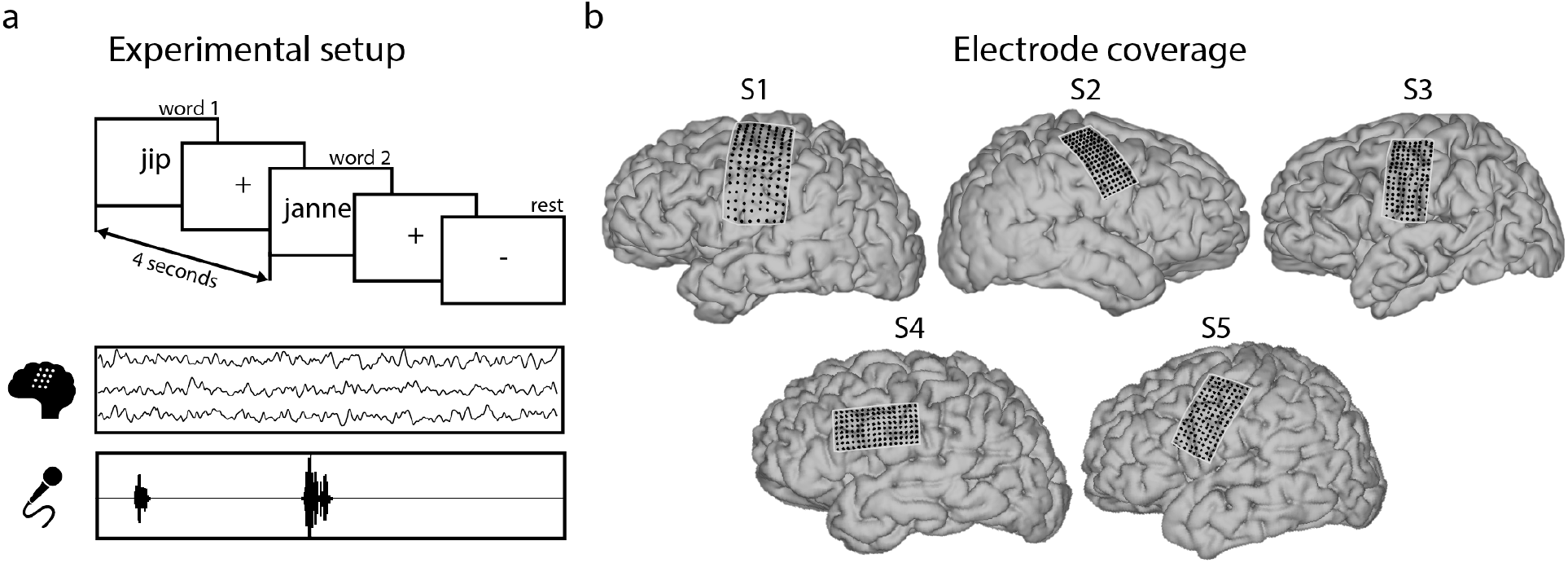
a. Experimental setup. b. Electrode coverage in all participants.

**Figure 2.**
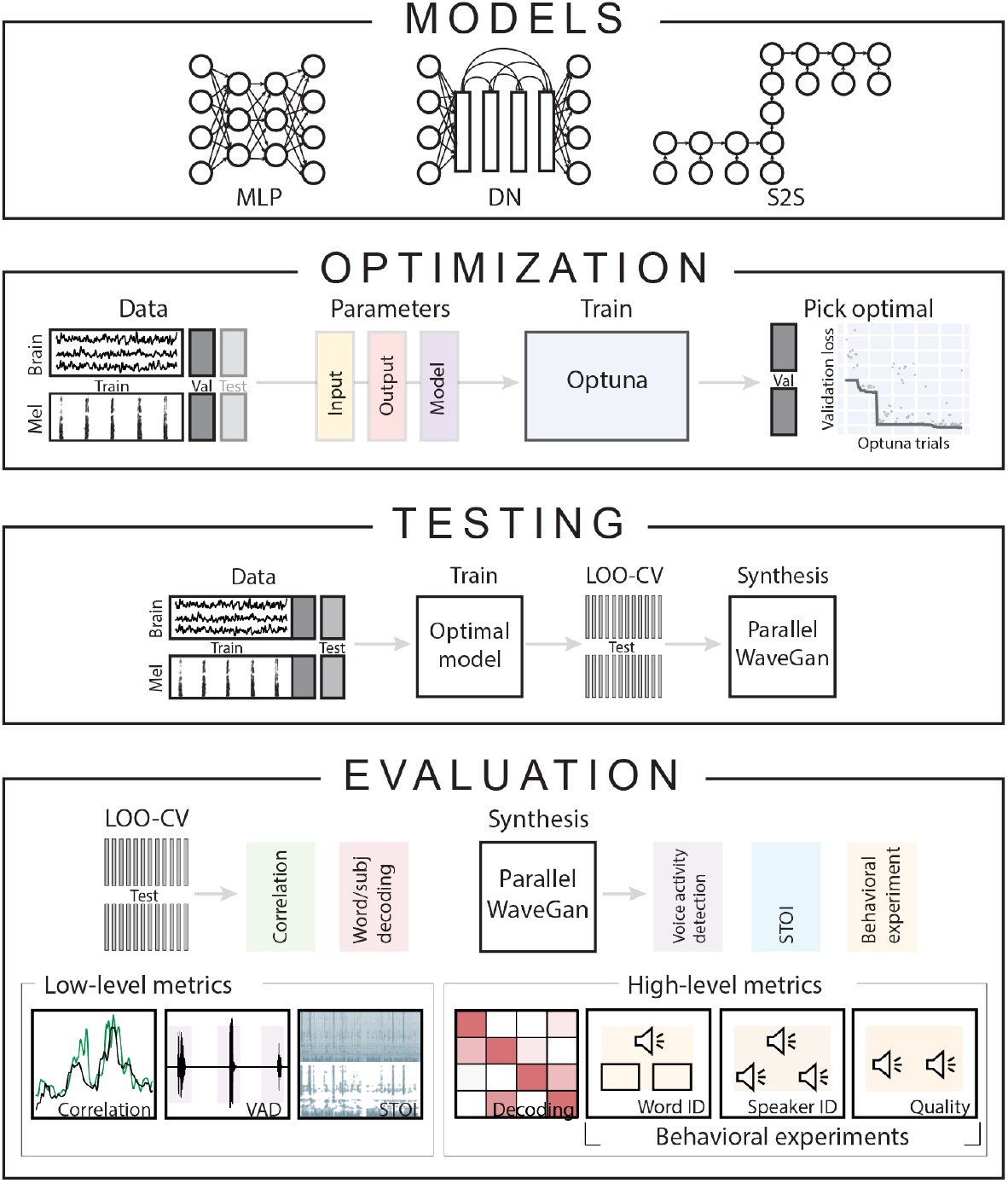
Overview of the approach. The present study trained, optimized and evaluated three deep learning speech reconstruction models: multi-layered perceptron (MLP), densenet (DN) and sequence-to-sequence (S2S). Each model was used to reconstruct target speech spetrograms based on input HD ECoG neural activity. Per HD ECoG subject, each model was optimized separately. For this, the task data was divided into train, validation and test sets. During optimization, input, output and model parameters were optimized in a way that minimized the speech reconstruction loss on the validation set. Once optimal set of parameters was identified, each model was retrained and tested on a held-out test set using leave-one-out cross-validation (LOO-CV). For speech synthesis from reconstructed and target spectrograms, a separate pretrained vocoder called WaveGan was used. For evaluation, we employed low-level metrics such as Pearson correlation, voice activity detection match (VAD match) and short-term objective intelligibility (STOI); and high-level metrics, such as word and speaker decoding with machine learning classifiers and perceptual judgments with behavioral experiments.

## Results

For speech reconstruction from sensorimotor brain activity we optimized and evaluated three widely-used artificial neural network architectures: a multi-layered perceptron (MLP), a densenet convolutional neural network (DN)^54^ and a sequence-to-sequence recurrent neural network (S2S)^55^. First, we report results of the optimization procedures (how successful they were and what parameters of the reconstruction model were optimized). Next, we report low-level metrics for evaluation of the reconstruction accuracy (Pearson correlation, match in voice activity detection, short-term objective intelligibility) and high-level metrics (identifiability of words and speakers with machine learning classifiers and speech perceptual intelligibility with human perceptual judgments). Finally, to determine which neural sources were most informative for achieving best results, we assessed the contribution of individual intracranial electrodes to the reconstruction accuracy.

### Model Optimization

#### Optimization of the reconstruction loss in validation and test data

The three model architectures (MLP, DN and S2S) were optimized separately due to differences in their hyperparameters. Each model was optimized per subject using the Optuna (https://optuna.org) framework^56^. We used a form of Bayesian optimization that constructed a distribution of the reconstruction loss given various parameter choices by taking into account the history of parameter changes (see Methods for more details). in, task data was divided into train, validation and test subsets. During training, the models minimized the reconstruction loss on the test set, measured as a mean squared error between target and reconstructed speech spectrograms. The reconstruction loss computed on the held-out validation set was used to optimize model parameters. After the optimization was complete, the reconstruction loss was computed on a separate test set to evaluate the results.

Overall, we observed a considerable drop in the reconstruction loss on the validation set, which consisted of a single trial of every unique word (Figure 3a). We also report substantial differences across subjects with respect to the minimal loss achieved (S1 and S5 reached much lower overall loss, especially in DN in S2S), and apparent interaction between subjects and models (S1 and S5 showed more distinct loss profiles across models compared to S2, S3 and S4, who showed similar profiles for MLP and S2S and whose minimal loss in DN and S2S was greater compared to S1 and S5).

**Figure 3.**
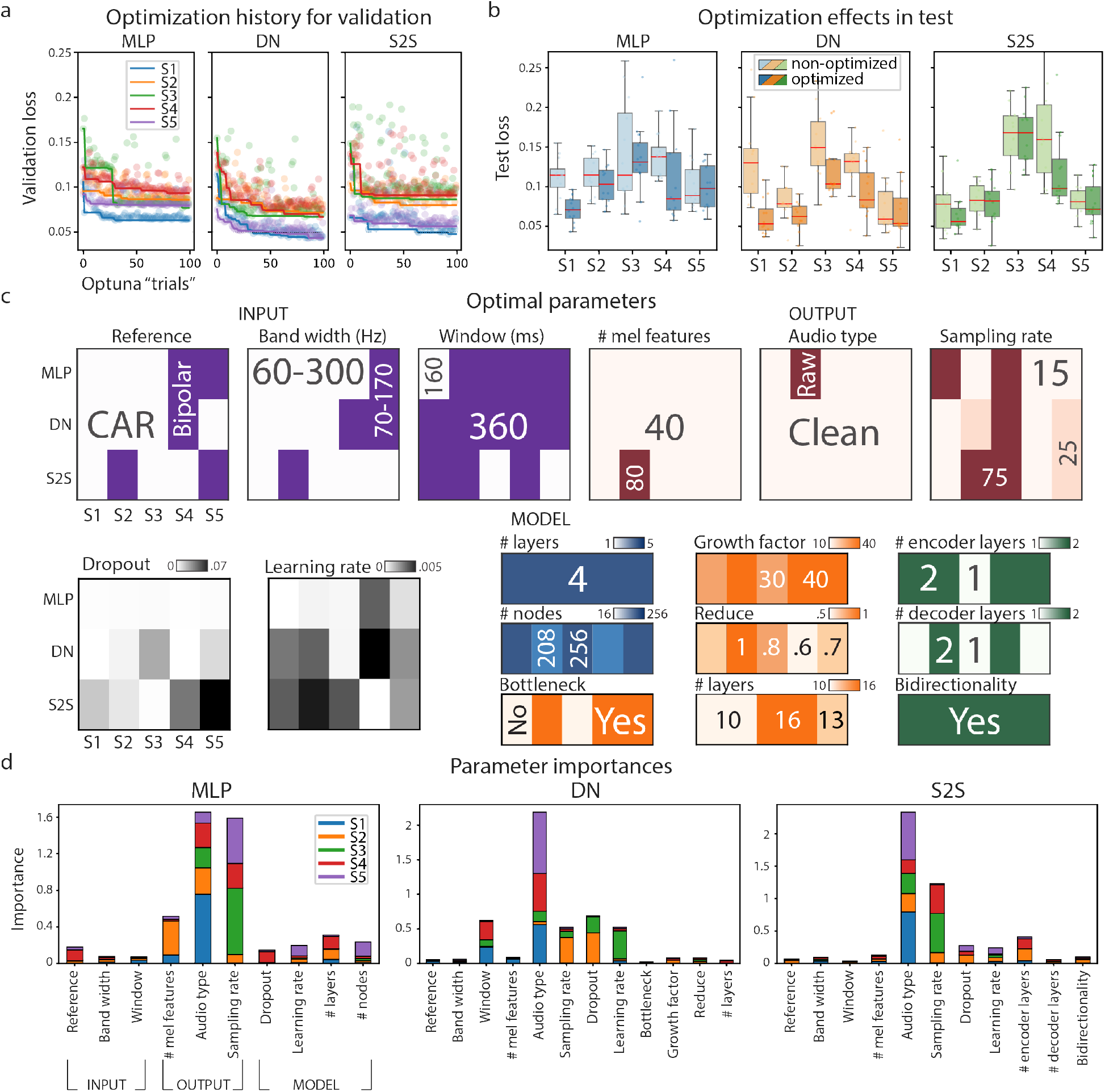
a. Optimization history for the validation set plotted over the course of the optimization procedure, i.e. Optuna optimization “trials” (see Methods for details). b. Optimization effects in the test set. Reconstruction loss computed on the test set is shown for non-optimized (a parameter set chosen at random at the start of the optimization procedure, i.e. first Optuna “trial”) and optimized models. c. Optimal parameters per model architecture (MLP, DN, S2S) and subject. Different colors represent different types of parameters: input (purple), output (red), general model hyperparameters, such as dropout and learning rate (grey) and model-specific parameters: MLP (blue), DN (orange) and S2S (green). These plots represent values of each parameter in the optimal model. For example, for S1 MLP best parameters were: CAR, 60–300 Hz, 160 ms window – input; 40 features of clean audio at 75Hz – output; dropout of 2*e* − 4, learning rate of 1*e* − 4, four layers of 256 nodes each. d. Parameter importances per model. Importance scores were normalized across parameters and are shown per model-subject pair.

Because the validation loss was used to guide model optimization (see Methods for details), it was possible that optimal models became overfitted to the validation data and therefore would not generalize to a held-out test set. We tested this possibility with a Wilcoxon signed-rank test in all subjects using loss values in a test set obtained with non-optimized and optimized models (Figure 3b). Non-optimized models used a parameter set that was chosen at random at the first step of the optimization process. The result showed a significant decrease of test loss in optimized models compared to non-optimized ones: *Z*_*all*_ = 5.9, *p* = 2 × 10^−9^), and separately per model: *Z*_*MLP*_ = 3.15, *p* = 6 × 10^−4^, *Z*_*DN*_ = 5.01, *p* = 3 × 10^−7^ and *Z*_*S*2*S*_ = 1.93, *p* = .03. This means that the observed optimization effects generalized to unseen test data that had not been part of optimization.

#### Optimal parameters and parameter importance

Since optimization of speech reconstruction models overall led to a decrease of reconstruction loss in both validation and test data, we asked next what were the optimal configurations of parameters associated with optimal models. The optimization routine tuned three types of parameters: 1) input (brain data), 2) output (target mel-spectrogram), and 3) model-specific parameters. All parameters and their ranges were selected based on previous literature and own experience. Parameters of the brain data inputs included the type of reference for de-noising HD ECoG signals (common average reference [CAR] or bipolar), band width of the frequency-transformed HD ECoG signal in the high gamma range (60-300 or 70-170 Hz) and temporal window of data used for modeling one spectrogram timepoint (160 or 360 ms). As target audio outputs for reconstruction we chose mel-spectrograms of speech, as they represent sound acoustics on a spectrum inspired by human auditory perception^57^. Parameters of the audio outputs reflected and type of audio preprocessing, as well as varied the details of calculating mel-spectrograms, and included the type of audio: microphone audio as it was acquired (raw) or de-noised microphone audio that filtered out all acoustics that were not produced by the subject speaking (clean, see Methods for details), the number of mel-frequency bins (40 or 80) and the sampling rate of the resulting spectrogram based on the choice of the window length and hop size parameters for spectrogram calculation (15, 25 or 75 Hz, see Methods for details). Finally, model-specific parameters included common for all models (learning rate and dropout) and architecture-specific parameters (number of nodes and number of layers in MLP; bottleneck, growth and reduce factor, number of blocks in DN, number of encoding and decoding layers and bidirectionality in S2S).

To explore optimal parameters, first we visualized optimal parameter configurations for each model-subject pair and each type of parameter (Figure 3c). Inspection of these plots allowed us to spot consistencies across models and subjects, such as dominance of CAR, 60–300 Hz of high gamma range and 360 ms temporal window among the optimal configurations of the input; preference for 40 mel features in clean audio at lower sampling rates (15 or 25 Hz) in the optimal configurations of the output. Across all model architectures there appeared to be a preference for a larger number of layers and nodes. In the case of S2S, optimal models in all subjects included bidirectionality.

To examine the effects that different parameters had on the loss values we used parameter importance estimates provided within Optuna. Overall, across subjects we observed that the reconstruction loss was most affected by output parameters, such as type of audio and sampling rate (Figure 3d). For MLP, the number of mel features and number of layers were the next most important parameters. For DN, the size of the input temporal window, dropout and learning rate were the next most important parameters. For S2S, the number of encoding layers, learning rate and dropout were the next most important parameters.

These results indicate that optimal configurations of parameters lead to better reconstruction losses. Specifics of audio processing may be of particular importance. Optimal parameters were often consistent across models and subjects. Yet it also appeared that some parameters may be subject and model-specific, highlighting the need for subject- and model-specific optimization for achieving best reconstruction results.

### Evaluation with low-level measures

Having reached optimal model performance in terms of the reconstruction loss, next we sought to evaluate the reconstruction quality. After brief visual inspection of the reconstructed and target spectrograms (Figure 4a) we turned to objective metrics of reconstruction quality typically used in the literature^19–21, 26^. These are measures based on low-level acoustic features that are easy to calculate and interpret. Three metrics were considered: 1) Pearson correlation between reconstructed and target speech fragments, 2) match in voice activity detection (VAD) between reconstructed and target fragments, and 3) short-term objective intelligibility measure (STOI)^58^. VAD match calculates the correspondence in detected voice activity between reconstructed and target waveforms. STOI, originally developed for comparison of clean and degraded signals, is an estimate of speech intelligibility based on the spectrotemporal feature correspondence between the audio fragments. In addition, we provided lower (based on random shifts) and upper (based on target audio fragments) bounds of model performance. We also focused on two comparisons of metric values: 1) between optimized and non-optimized models (within-model), and 2) across the three optimized models (MLP, DN and S2S).

**Figure 4.**
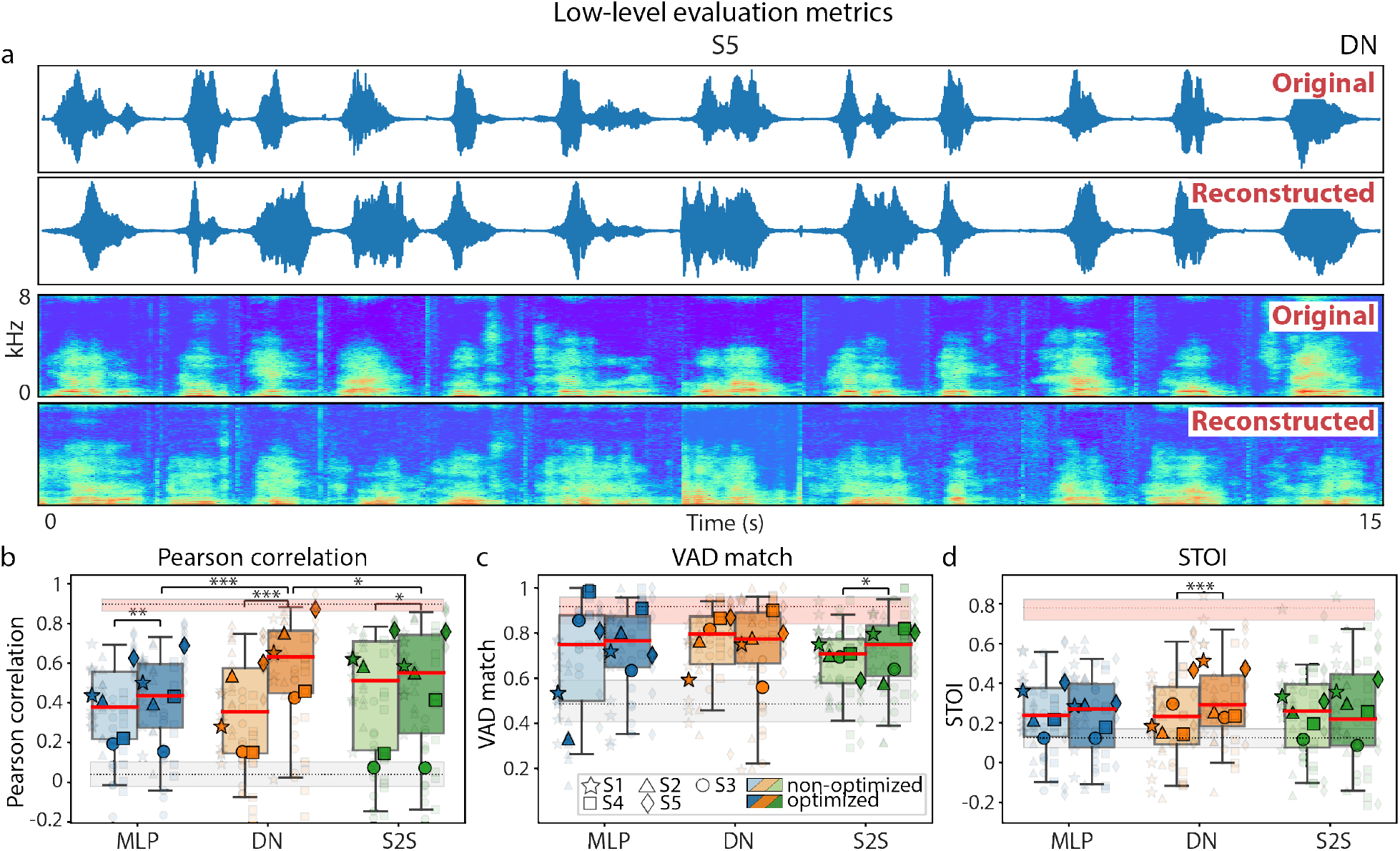
a. Examples of original and reconstructed spectrograms (test data, DN model, S5). b-d. Low-level evaluation metrics computed on test data and shown for non-optimized and optimized versions of each model: Pearson correlation (b), voice activity detection (VAD) match (c), short-term objective intelligibility (STOI, d). Boxplots outline 25th and 75th percentile of results across test trials of all subjects. Red line shows medians. Semi-transparent smaller markers show results per individual test trial of each subject. Opaque bigger markers show averages across test trials per subject. Lower (based on data shifts) and upper (based on target audio waveforms) performance bounds are shown in grey and pink, respectively. Results for the three model architectures are shown in different colors (MLP in blue, DN in orange and S2S in green). Color saturation denotes the use of optimization: results for non-optimized models are shown as non-saturated and results of optimized models are saturated. Significant differences in medians are marked: *p* < .05 (*), *p* < .01 (**) and *p* < .001 (***).

Pearson correlations, calculated between reconstructed and target spectrograms, revealed the largest optimization effects compared to VAD match and STOI (Figure 4). The difference between optimized and non-optimized reconstruction models in terms of Pearson correlation scores was significant for all models: *Z*_*MLP*_ = 2.67, *p* = 4 × 10^−3^, *Z*_*DN*_ = 5.85, *p* = 2 × 10^−9^ and *Z*_*S*2*S*_ = 1.66, *p* = .05. This was somewhat expected since Pearson correlations are related to the reconstruction loss (measured as the mean squared error between predictions and targets). In addition, Pearson correlations in DN were significantly greater compared to MLP and S2S: *H* = 14.26, *p* = 8×10^−4^, based on a non-parametric Kruskal-Wallis test and post-hoc Dunn test on individual comparisons: *p*_*DN− MLP*_ = 1 × 10^−4^ and *p*_*DN− S*2*S*_ = .04.

VAD match and STOI were calculated on synthesized waveforms. This allowed us to provide ceiling estimates for each subject-model pair by computing the metric value on original audio fragments (used as input to the reconstruction models) and audio fragments resynthesized from spectrograms to waveform using the external speech synthesis model (Parallel WaveGAN vocoder). Both VAD match and STOI showed smaller effects of model optimization: *Z*_*S*2*S*_ = 1.87, *p* = .03 (for VAD) and *Z*_*DN*_ = 3.27, *p* = 5 × 10^−4^ (for STOI), and no significant effect of model architecture. Overall, VAD match median (across subjects) scores were closer to the upper bound performance than median STOI scores, which despite being significant in most cases, were quite low. STOI was the only metric for which individual subject results did not reach significance in any of the models (S3 and S4).

Similar to the results from the section on Model Optimization, we also observed considerable variance in metric scores across five subjects with S1, S2 and S5 on average obtaining greater Pearson correlation and STOI scores compared to S3 and S4.

It is important to note that these metrics rely on low-level acoustic features and are convenient to consider when evaluating audio reconstruction quality (especially given the low-level spectrogram feature loss used in training). In the case of speech and the ultimate goal of using these reconstruction models in clinical BCI applications, it makes sense to look beyond low-level quality metrics and assess reconstruction of high-level characteristics of speech.

### Evaluation with high-level measures

Therefore, next we analyzed high-level properties of the reconstructions, such as identifiability of individual word and speakers, as well as intelligibility and perceptual quality of resulting speech. First, we assessed word and speaker identifiability with objective metrics, such as machine learning classifiers. Then, in two separate speech perception experiments, we collected subjective human judgments of word intelligibility and speaker recognition. Finally, in a third behavioral experiment, we collected and analyzed perceptual quality judgments of the reconstructions.

#### Machine learning classifiers

We trained a linear logistic regression classifier to decode word identity (out of twelve individual words). For audio reconstructions, a linear logistic regression classifier was trained on target audio spectrograms per subject and tested on corresponding reconstructions from the test set per subject and model (including optimized and non-optimized versions of each model). Word classification accuracy was highly significant (reaching accuracy of 92-100%, chance level of 8%) across all subjects and models. There was no significant difference in accuracy between optimized and non-optimized models, different model architectures or subjects (5a).

Given the observed result, next, we asked ourselves whether using audio reconstructions for word classification would provide any boost in accuracy compared to the use of raw brain activity (that was fed into the reconstruction models) as input to the classifier. To test this, we retrained each logistic regression on brain input rather than audio spectrograms and tested it on the same test set of 12 words as above. Interestingly, we observed a much larger variance in word classification accuracy across subjects compared to the use of audio spectrogram reconstructions (Figure 5a), consistent with inter-subject variability patters from low-level evaluation metrics (lower performance of S3 and S4). The median accuracy across subjects was greater for the classifiers trained on reconstructions: *Z*_*MLP*_ = 6.51, *p* = 4 × 10^−11^, *Z*_*DN*_ = 6.73, *p* = 9 × 10^−12^ and *Z*_*S*2*S*_ = 6.73, *p* = 9 × 10^−12^.

**Figure 5.**
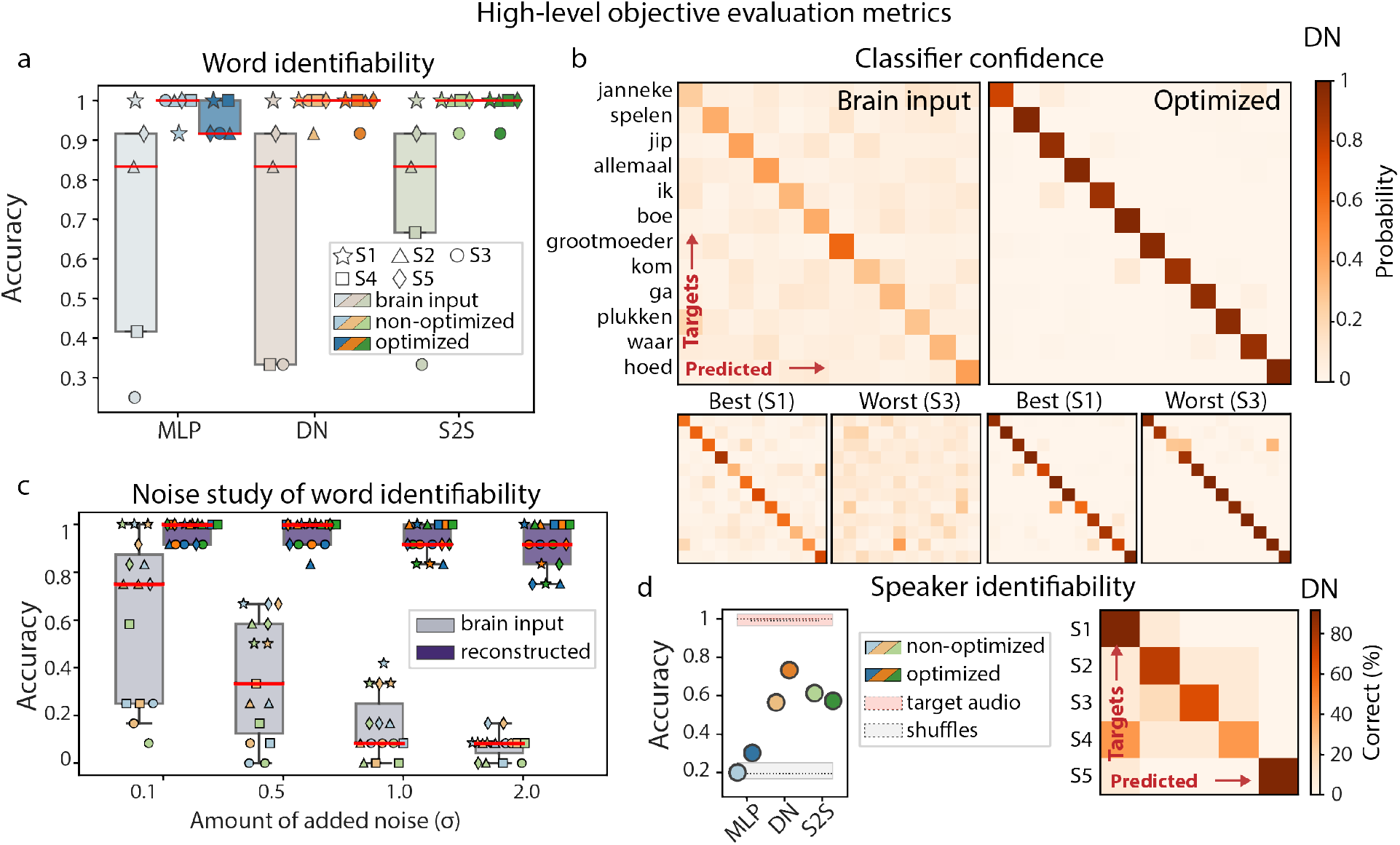
a. Word identifiability as assessed with word classification using audio reconstructions from optimized and non-optimized versions of the three models (MLP, DN, S2S) as well as the raw brain input. B. Probability matrices for predicted and target test words for a classifier trained on brain input, and for a classifier trained on audio spectrograms (tested on reconstructions). Top panel shows probability matrices averaged over subjects. Optimized DN model was used to obtain the reconstructions. The lower panels represent matrices of best (S1) and worst performing (S2) subjects with respect to the word classification accuracy. The matrices are shown per type of input: brain input and audio reconstruction with optimized models. c. Results of the noise study: performance of the classifiers trained and tested on noisy inputs. The amount of noise was gradually increased. Individual markers show performance per subject. The three model architectures are shown in different colors (MLP in blue, DN in orange and S2S in green). Color saturation denotes the type of input data: brain input (least saturated), speech reconstructions obtained with non-optimized models (medium saturated) and speech reconstructions obtained with optimized models (most saturated). For each optimized speech reconstruction model, either resulting reconstructions or raw brain data (to be passed as input to the speech reconstruction model) was used an input to the machine learning classifier. Therefore, in the case of brain inputs, brain data with optimal input parameters per model (sampling rate, reference and HFB range) was used. d. Speaker identifiability as assessed with speaker classification using classifiers trained on audio spectrograms and tested on reconstructions. Results for optimized and non-optimized versions of the three models (MLP, DN, S2S) are shown. A confusion matrix for the best performing classifier (optimized DN) is shown on the right.

In addition, examining classifiers’ confidence about the predicted word classes (expressed in probability distributions over all candidate classes per trial) showed considerably lower confidence in classifiers trained on brain input compared to audio reconstructions (Figure 5b) with median confidence of .3 and 1, respectively. It appeared that classifiers trained on audio spectrograms were more robust across all subjects compared to classifiers trained on brain input. To test this further, we performed a noise study, in which we repeatedly added white noise (from standard normal distribution) to the input training and test data of the classifier, retrained it and recalculated its accuracy on the test data. We observed that word identification accuracy changed as a function of amount of noise and decayed rapidly when the classifier used brain input. Using audio reconstructions, on the other hand, resulted in noticeably more robust classification accuracy, even when considerable amounts of noise were added (5c).

Next, we trained logistic regression classifiers to decode speaker identity. As with word classification, the classifiers were trained on target spectrograms and tested on reconstructions for optimized and non-optimized versions of each model. We observed the highest accuracy for optimized DN model, reaching 73% (*p*_*DN*_ = 0 based on random label permutations, chance level is 20%, 5d). There was a trend for optimized models to perform better (in DN and MLP, but not S2S). DN and S2S models achieved higher accuracy compared to MLP. The latter provided insignificant decoding accuracy: *p*_*MLP*_ = .45, *p*_*MLP*_ = .2 for non-optimized and optimized models, respectively. Due to the difference in the number of channels in input brain data, no classifiers on brain input were trained for speaker decoding.

#### Human perceptual judgments

With two follow-up speech perception experiments we sought to test whether word and speaker recognition not only were successful with machine learning classifiers trained on target spectrograms, but were also possible for human perceptual judgments.

In experiments I and II, 30 and 29 healthy Dutch-native volunteers respectively, were presented with an audio reconstruction and two options, of which they were asked to choose one that suited the reconstruction best. In the word intelligibility experiment, the two options were two written words (for example, “janneke” and “grootmoeder”), one being the correct word, and the other chosen at random from remaining 11 words. In the speaker recognition experiment, two options were target audio fragments of two speakers pronouncing the same word as in the reconstruction (for example, “janneke” said by S1 and S4), one being the correct speaker, and the other chosen at random from remaining 4 speakers. Both experiments included catch trials that used target audio fragments instead of reconstructions. This was done to obtain ceiling estimates of word intelligibility and speaker recognition.

In the case of word intelligibility, participants on average were able to identify words significantly above chance across subjects and models (except for some models of S3 and all models of S4): mean accuracy values reached 59%, 61%, 65% for MLP, DN and S2S models, respectively. Best scores were obtained for DN and S2S models: *H* = 7.67, *p* = .02, based on a non-parametric Kruskal-Wallis test and post-hoc Dunn test on individual comparisons *p*_*S*2*S*_− _*MLP*_ = 7 × 10^−3^ and *p*_*S*2*S*_−_*DN*_ = .06. (Figure 6a). Moreover, there was a positive relationship between low-level Pearson correlation and high-level word intelligibility metric for DN and S2S models.

**Figure 6.**
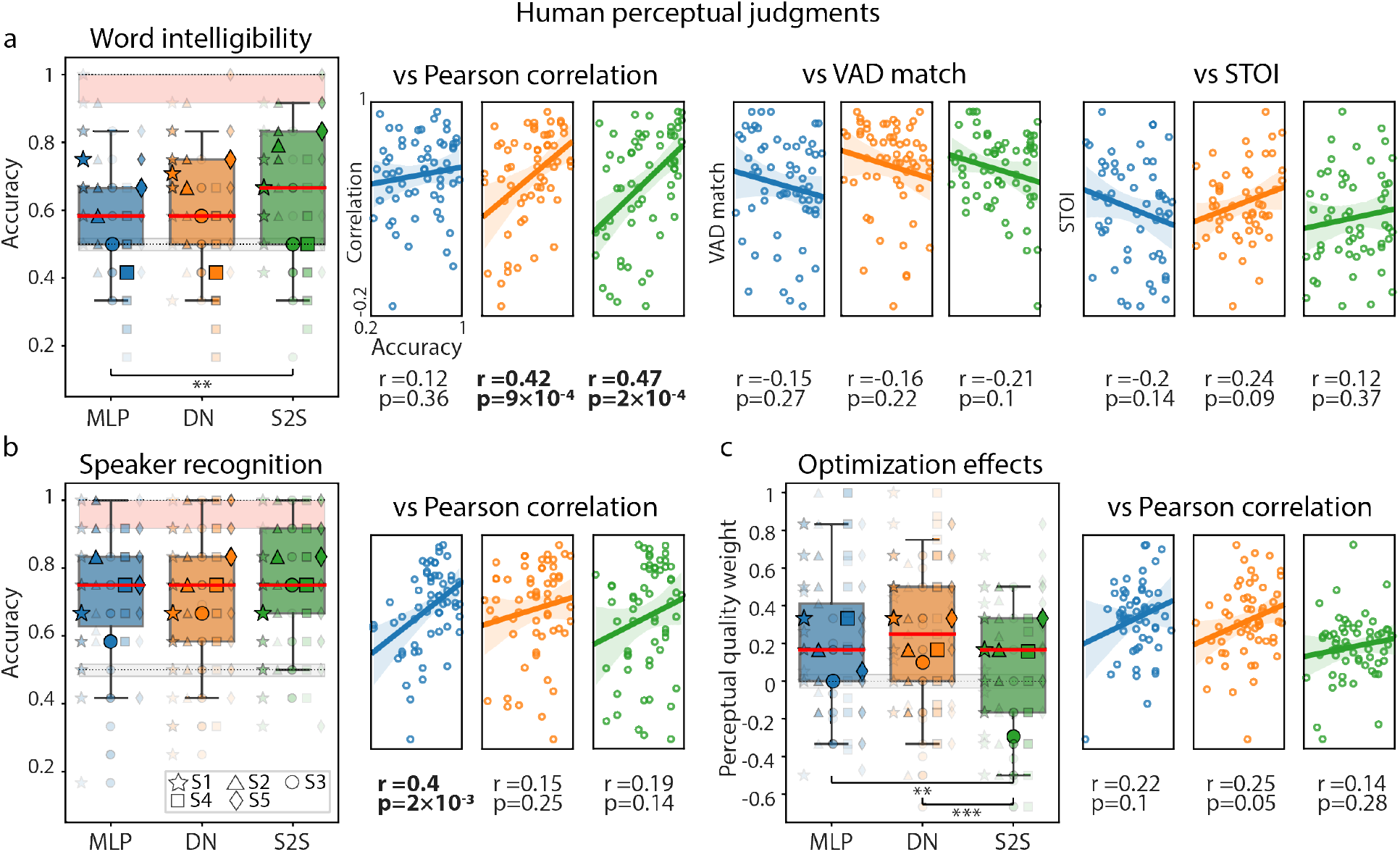
Human perceptual judgments obtained with three behavioral experiments. a. Word intelligibility (experiment I). b. Speaker recognition (experiment II). c. Perceptual quality of optimized and non-optimized model reconstructions (experiment III). Perceptual judgments per each experiment were plotted against low-level evaluation metrics (scatter plots to the right of each boxplot panel). Boxplots outline 25th and 75th percentile of behavioral results. Red line shows medians. Semi-transparent smaller markers show results per behavioral participant on data of each HD ECoG subject (averaged over 12 test word trials). Opaque bigger markers show averages across behavioral participants per HD ECoG subject. Results for the three model architectures are shown in different colors (MLP in blue, DN in orange and S2S in green). Lower (based on randomly assigned perceptual judgments) and upper (based on target audio waveforms) performance bounds are shown in grey and pink, respectively. Experiment III did not use target audio waveforms and therefore does not have the upper performance bound. Significant differences in medians are marked: *p* < .05 (*), *p* < .01 (**) and *p* < .001 (***). Boxplots in experiments I and II show accuracy of word and speaker recognition based on binary comparisons. Boxplots in experiment III show whether reconstructions from optimized or non-optimized models were judged to be better (more intelligible, less noisy). An arbitrary value (perceptual quality weight) of 1 was assigned to each judgment if the audio produced by the optimized model had been judged as better and a value of -1 otherwise (see Methods for more details). In order to make scatter plots of perceptual judgments against low-level metrics, perceptual judgments were recalculated per test word trial (averaged over behavioral participants). Therefore, each data point of a scatter plot represents data per single word from the test set of each HD EcoG subject – 60 data points in total (12 word trials × 5 subjects).

In the speaker recognition task, participants showed even better overall performance and a large gap between behavioral scores and permutation-based lower bound: mean accuracy values reached 72%, 73%, 75% for MLP, DN and S2S models, respectively. There was no significant difference in scores across models and the only model with a positive relationship between Pearson correlation and speaker recognition judgments was MLP (Figure 6b).

These results differ from the performance of machine learning classifiers for both word intelligibility and speaker recognition. In the case of word intelligibility, machine learning classifiers reached an accuracy of 92-100% (chance level 8%) in identifying individual words, whereas behavioral judgments indicated that perceptual quality of reconstructed words was considerably lower and provided an average accuracy of 58-66% (chance level 50%). In both cases, the MLP model was outperformed by DN and S2S. For speaker recognition, machine learning classifiers showed best results for DN and S2S as well: 73% and 57% respectively (chance level 20%), with behavioral judgments achieving 75% (chance level 50%) for all models. Due to the differences in the computation of accuracy between machine learning classifiers and perceptual judgments, direct comparison between the two results cannot be performed. Moreover, behavioral experiments were performed on synthesized speech waveforms whereas machine learning classifiers used normalized spectrogram data. In order to synthesize waveforms the spectrograms were re-scaled and centered with mean and standard deviation of the target audio and passed through the external vocoder. This was necessary for correct processing by the vocoder but may have affected perceptual quality and speech intelligibility.

In another behavioral experiment (experiment III) we sought to assess model optimization effects on perceptual scores. Similar to the previous experiments, 29 participants were presented with two options: non-optimized and optimized reconstruction audio fragments, otherwise matching in word, speaker and model (for example, optimized and non-optimized “janneke” reconstruction using the MLP model on data of S1). No catch trials using target audio were used. We found that, on average, participants judged reconstructions from optimized models to be perceptually better across all models. Largest perceptual gains were obtained for optimized DN and MLP models: *H* = 18.74, *p* = 8 × 10^−5^, based on a non-parametric Kruskal-Wallis test and post-hoc Dunn test on individual comparisons *p*_*DN*− *S*2*S*_ = 3 × 10^−5^ and *p*_*MLP* − *S*2*S*_ = 3 × 10^−3^. (Figure 6c). This result confirmed not only that optimization led to a lower reconstruction loss and more accurately preserved low-level features (measured with Pearson correlation), but also that optimized models were associated with better perceptual quality of the reconstructions.

Altogether, our behavioral results further supported the previous conclusions made based on the loss analysis, low-level and objective high-level evaluation metrics, and confirmed overall high-quality of the optimized speech reconstructions.

### Contribution of individual electrodes

Finally, given overall high-quality reconstruction accuracy, as assessed with multiple low-level and high-level metrics, we sought to investigate the relationship between model performance and input brain features. Understanding interactions between electrode location on the sensorimotor cortex and its influence on speech reconstruction quality is vital for future applications of this work in the BCI field. To assess this, we performed an input perturbation analysis to assess individual contribution of each HD ECoG electrode to the reconstruction accuracy. For this, during testing (after models had been normally trained) per electrode we perturbed brain data that was input to the reconstruction models (optimized MLP, DN or S2S) by replacing that electrode’s time course with zero values (done after centering the input data) and recording the resulting loss value. Thus we were able to associate perturbation of data per electrode with the amount of change in the loss value compared to using all input data.

Such input perturbation analysis revealed that most informative electrodes (those associated with largest increases in the loss value, and consequently, worse reconstruction accuracy) were grouped in small clusters along the ventral and dorsal premotor and motor areas, rather than being evenly distributed across the HD ECoG grids (Figure 7). We also observed a degree of variability across patients and, to a smaller extent, across reconstruction models. This, together with previously mentioned differences across subjects may indicate that different cortical locations can be more or less informative for speech decoding across subjects, and that an individual subject approach is beneficial for achieving high accuracy of results.

**Figure 7.**
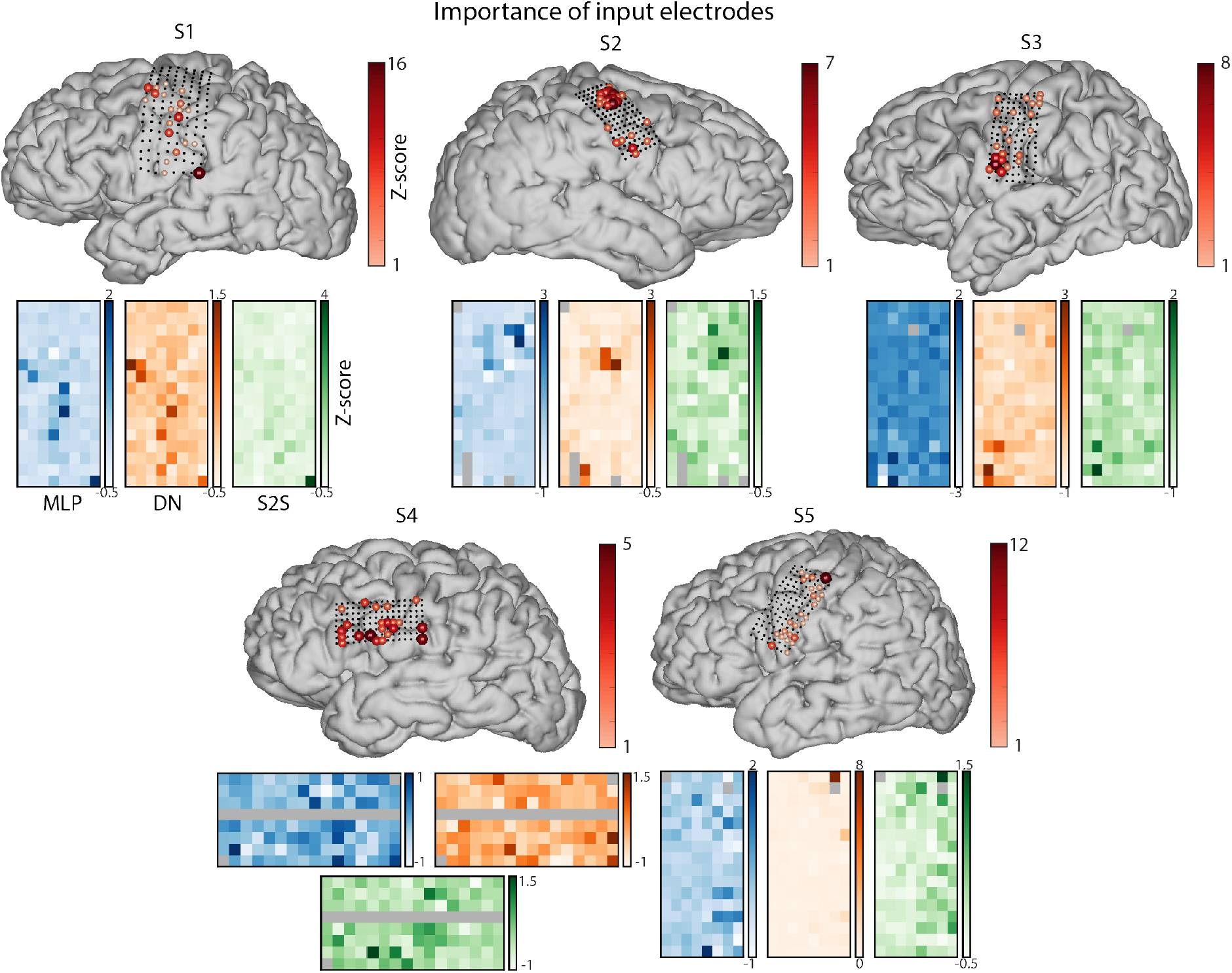
Results of the input perturbation analysis per electrode showing how much the reconstruction loss increases depending on perturbation of individual electrodes. Brain points show a sum of loss increases over all models. Grid plots below show z-scored loss changes per model (positive: loss increase, negative: loss decrease compared to the baseline of using all electrodes). Results for the three model architectures are shown in different colors (MLP in blue, DN in orange and S2S in green). HD ECoG electrodes excluded from all analyses due to bad signal quality are shown in grey.

## Discussion

In this study, we performed a systematic optimization and evaluation of models for speech reconstruction directly from sensorimotor brain activity recorded with HD ECoG grids. We found that end-to-end deep learning approaches, currently dominating the field, overall benefited from model optimization, and that the choice of output parameters of these models (target spectrograms) had the largest effect on the reconstruction quality. Next, we showed that optimized models led to overall high reconstruction quality as assessed with both low-level evaluation metrics and high-level speech measures, such as word and speaker identifiability and perceptual quality. Word recognition in reconstructed audio was markedly more accurate, stable across subjects and robust compared to raw brain input. Behavioral experiments confirmed individual word intelligibility and optimization gains in perceptual judgments. Finally, we quantified the relationship between the reconstruction accuracy and location of HD ECoG electrodes, revealing that the largest contribution to model performance was made by small clusters of electrodes throughout the ventral and dorsal premotor and motor cortices. These results have the potential to further advance the state-of-the-art in speech decoding and reconstruction in severely paralyzed patients who rely on BCI technology for communication.

### Optimized complex model architectures lead to best reconstruction of spoken speech from brain signals

For all models, optimization led to a decrease in the reconstruction error. Moreover, behavioral quality judgments confirmed that in all models optimization led to an overall higher perceptual speech quality. Importantly, in this study we optimized input, output and model-specific parameters of speech reconstruction models. Our results indicate that input parameters, or options for preprocessing brain data, and model-specific parameters may have had a lower effect on performance compared to the output parameters, or the choices of target audio features. Next to the use of coarser spectrograms (40 frequency bins at 15-25 Hz), we demonstrate that quality of microphone recordings and processing steps aimed at cleaning noisy signals are key for achieving best performance. This appears logical, but the clinical setting in which ECoG data are often acquired (especially for experiments with participants during awake surgery) can be a challenging environment for collecting high-quality data, and this needs be taken into account when planning and conducting the research. Moreover, this result may contribute to the discussion about the type of information that drives neural responses. Background noise removed from “clean” audio signals was part of the auditory input to the brain, yet in most subject-model pairs, included S1 who had limited auditory coverage, reconstruction was best if only produced speech, and not all perceived sound, was used.

After optimization, we compared the performance of the deep learning architectures used in speech reconstruction. Three model architectures, popular in the field of computational neuroscience and neuroengineering, were considered: a simple multi-layered perceptron (MLP) model only consisting of linear layers followed by a non-linear activation function, and two more complex models: a convolutional DN model, and a recurrent sequence-to-sequence (S2S) model (see Methods for details). All these architectures have previously been employed in studies on speech decoding with limited attempts to optimize and compare results across model architectures^19, 20, 26^. We observed that more complex models, such as DN and S2S, outperformed MLP in a number of comparisons: larger median Pearson correlation (DN), higher median word and speaker classification accuracy (DN and S2S) and better perceptual judgments of word identifiability (S2S). Moreover, among various MLP hyperparameter configurations, those with a larger number of MLP layers (4 from the range of 1 to 5 layers) and nodes (208 and 256 from the range of 16 to 256 nodes) turned out to generate the best results for this approach. These results suggested that high complexity of deep neural network models is required to achieve more accurate speech reconstructions. Our results also indicate larger optimization effects in DN and MLP compared to S2S (Figure 3b, 4a, 5c). And even though DN occasionally outperformed S2S, the improvements were marginal, which makes it difficult to conclude that convolutional architectures should be preferred over recurrent ones.

Recent work in computational neuroscience suggests that the use of recurrent connections and attention modules can be beneficial in processing brain signals^59–61^. Recurrence can help capture local and long-range temporal dependencies in the data whereas attention helps to effectively process temporally organized inputs. It is important to note that the present results have been obtained with relatively small datasets, in which subjects pronounced individual words. It is possible that for speech data that exhibits longer temporal dependencies, such as speech phrases, sentences and narratives, attention-based encoder-decoder models will further outperform convolution-only architectures, such as DNs^26^.

### Individual evaluation metrics highlight different aspects of model performance

Evaluation of the speech reconstructed from sensorimotor cortex was performed using various low- and high-level metrics. We observed that each metric assessed the model performance differently. Pearson correlation metric showed the largest improvement of the reconstructed speech over the permutation-based baseline and was the only low-level metric to reflect model optimization gains. In general, this result appears logical as there is a straightforward relationship between the reconstruction loss (mean squared error) and Pearson correlation of predictions and targets. In the future, more sophisticated loss functions may be explored especially for achieving higher perceptual quality of the reconstructions, which could be regarded as the ultimate test of performance in BCI applications for restoring communication. VAD match metric showed improvement compared to the baseline, yet no model optimization effects. It is possible that distinguishing between reconstructed speech and silence may have been a simple enough task that did not require optimization. Finally, due to the lack of significance and overall low values, STOI metric did not seem to be informative above and beyond what the other two measures showed. This could in part be due to the way it is calculated in that it requires the audio fragments to be at least 300 ms in length. This limitation can be particularly inconvenient when reconstructing individual monosyllabic words. Altogether, low-level assessment of reconstruction results indicated that Pearson correlation remains the simplest and most straightforward low-level metric for evaluation of reconstruction accuracy in individual words, as long as the mean-squared error, otherwise known as the “pixel loss” is used for model training. VAD and VAD match can remain useful in identifying speech fragments in the brain signal, especially if such identification is not trivial, such as during covert or attempted speech. STOI as an evaluation metric may potentially be phased out.

The present study also explored several high-level metrics. Machine learning classifiers were employed to assess speech identifiability, and human perceptual judgments were collected to rate intelligibility and perceptual quality of reconstructed speech. Machine learning classifiers demonstrated high levels of word and speaker identifiability in reconstructed speech. Perceptual judgments of speech intelligibility seemed to agree to some extent with Pearson correlation and machine learning classifiers. It was clear, however, that perceptual quality was a unique measure that could not fully be captured in any other objective measure considered here.

### Direct speech reconstruction from sensorimotor cortex as the basis for next-generation BCI applications

Importantly, the results of this study indicate that speech reconstruction from HD ECoG data could be used for word decoding that is superior to decoding directly from raw brain activity. We have shown that reconstruction-based word decoding can reach 100% accuracy of decoding twelve words if complex models, such as DN and S2S, are used (Figure 5a). It is highly stable across subjects and can therefore reduce subject-specific variability (Figure 5a) and it is robust against noise, as classifier probability distributions over all possible word classes greatly favor the target of decoding (Figure 5b, c).

It is not trivial to directly compare these results with other word decoding work from sensorimotor cortex in the BCI field. However, to our knowledge, existing studies so far have been able to reach accurate word classification results during speech production by decoding from raw ECoG brain input (average accuracy of 85 ± 13% on 45 binary word classification tasks^2^ and average accuracy of 47.1% of decoding 50 words^33^) and on average used more repetitions of each word (30 repetitions of 10 words^2^ and 34 repetitions of 50 words^33^) compared to the present study (10 repetitions of 12 words). Our word decoding results using speech reconstructions from HD ECoG data show unprecedentedly high accuracy and robustness of decoding on a relatively small dataset and highlight the potential of this approach for further use in BCI.

Training word decoders on external speech data (for example, publicly available large audio corpora) and testing them on reconstructions from ECoG data from unseen subjects is the next step towards future application of this work in BCI. Highly accurate (>90% accuracy) and reliable discrete decoding of several classes of words (compared to the state-of-the-art binary decoding^31^) will help BCI devices offer more degrees of freedom to their users and thereby provide a new state of the art in the field. Moreover, further increasing the number of decodable words to a set of 50 or 100 commonly used words can lead to the development of the next-generation BCI technology for communication^33^.

Making the strategy of speech decoding though direct reconstruction from neural data work in a BCI setting with paralyzed individuals will pose some challenges. For training the reconstruction model we need targets of decoding associated with neural activity. This can be overcome by using speech sounds as cues and training paralyzed participants to mimic speech they heard. Then, speech cues can be used as targets. Alternatively, transfer models^62, 63^ need to be considered. Such models can potentially be trained on able-bodied participants and learn speaker-invariant audio representations and use external sound generation models^64^ to aid speech reconstruction.

### Informative neural sources may be distributed across sensorimotor cortex

Apart from the neural targets and decoding model, the results of the present study also contribute to the discussion about the neural sources of decoding and the details of the acquisition technique. Specifically, the present study showed successful speech reconstruction and word identifiability using HD ECoG electrodes over the sensorimotor cortex – one of the most promising regions in BCI implants for communication. It is important to note that other regions, such as Heschl’s gyrus and superior temporal cortex, were not the focus of this study, as their primary function is auditory processing of perceived speech. In the present work, we used a technique called input perturbation analysis to identify HD ECoG electrodes that were most informative for speech reconstruction and word decoding. Our results indicate that multiple regions throughout the motor, premotor and potentially inferior frontal gyrus contribute to accurate speech reconstruction. This is largely in line with previous work^4, 9, 10, 15^. Interestingly, subjects with best reconstruction results (S1, S2 and S5) benefited from contributions of electrodes located in the dorsal premotor and motor cortex (Figure 7). The subject who did not have dorsal motor coverage (S4) showed contribution of electrodes in the inferior frontal gyrus, but their contribution scores were lowest compared to those of other subjects, who had electrodes located in dorsal premotor and motor cortex. These results may indicate the potential of dorsal premotor and motor regions in speech reconstruction and are in line with the previous reports on speech decoding from that region^25, 28, 33^.

### Inter-subject variability of results and concerns for BCI

We also observed that despite optimizing individual subject’s datasets, there was a large variance in model performance across subjects. This inter-subject variability is a common outcome in speech decoding studies^12, 20, 23^ and can stem from several factors. One key factor is the choice of the brain area that is covered with electrodes. Missing an important patch of cortex can dramatically reduce performance. Electrode placement, however, is influenced by clinical considerations such as orientation of the grids to accommodate the leads, anchor veins that make optimal positioning impossible, and lack of knowledge on which cortical patches are most informative. Other factors include patient alertness and motivation, medication and epileptic spikes that confound the brain signals. In the present study, data from S4 was collected during awake surgery, where participant alertness may have been considerably reduced. Bigger datasets with larger numbers of participants would be beneficial for systematic investigation of these factors. Their optimization at data collection, processing and decoding stages may be key for further progress in the field.

Importantly, we also find that even though low-level features and perceptual quality of reconstructions varied across participants, word identifiability in reconstructed speech remained accurate and robust across all subjects. These results highlight the possibility of using direct speech reconstruction as a tool for boosting word decoding from brain data.

## Limitations

The present work has a number of limitations. Given a wide range of options and a lack of consensus regarding the effects of parameter choices (input, output and model-specific parameters) in the literature, here we limited ourselves to a few sets of parameters. Our choices were informed by existing literature and own experience, however, several improvements can be considered in the follow-up work. First, some recent work has demonstrated the potential of lower-frequency information and its coupling with the high-frequency band for speech processing and decoding^19, 28, 65^. Therefore, future work on speech reconstruction from sensorimotor cortex could explore the effects of adding lower-frequency ECoG components.

Second, several of the input and output parameters in the present work were considered to be categorical variables during optimization (for example, band width of the frequency-transformed signal in the high gamma range and temporal window of brain data used for modeling one spectrogram timepoint) with only a limited number of values used. This could be improved in the follow-up work in order to consider a more appropriate distribution over these parameter choices, include a larger range of options and achieve better optimization.

The present study implements an input perturbation analysis in order to identify electrode contribution to reconstruction accuracy. Our implementation of this method has a number of limitations. Specifically, it does not take into account multivariate activity that is likely to be informative for reconstruction and instead simplifies the spatial relationship between individual electrodes. Similar to the points made above, electrode contribution and selection could potentially be implemented as part of the optimization procedure as well.

## Conclusion

In this study, we performed dedicated optimization and evaluation of different deep learning models for speech reconstruction directly from intracranial sensorimotor neural activity during a speech production task. We showed that machine learning optimization of speech decoding pipelines and detailed evaluation of model performance metrics allow for achieving more accurate and interpretable reconstruction results, and improve our understanding of the brain signal and its relation to the speech features. Altogether, our results indicate that direct speech reconstruction from sensorimotor brain activity provides highly accurate and robust word decoding performance and overall intelligible speech, and therefore, has the potential to advance the state of the art in real-world applications of brain-computer interfaces for communication in severely paralysed individuals.

## Methods

### ECoG experiment

#### Participants

Four participants (S1, S2, S3, S5, age 36, 30, 51 and 24, respectively; three females) with medication-resistant epilepsy were admitted to the intensive epilepsy monitoring unit for diagnostic procedures after they underwent temporary implantation with subdural ECoG electrode grids to determine the source of seizures and test the possibility of surgical removal of the corresponding brain tissue. In addition to clinical procedures, participants gave written informed consent to participate in scientific research that accompanied ECoG recordings and could be conducted between clinical procedures. Participants also gave written informed consent for implantation of an additional high-density (HD) ECoG grid over their sensorimotor cortex for research purposes. One more participant (S4, age 21, male) underwent clinical awake surgery for brain tumor removal. During the surgery, a grid of HD ECoG electrodes was placed subdurally over the sensorimotor and inferior frontal cortices while the subject participated in functional mapping (up to 10 minutes) to guide tumor resection. The participant gave written informed consent to acquire and use their HD ECoG data from functional mapping tasks for research purposes. The study was approved by the Medical Ethical Committee of the University Medical Center Utrecht in accordance with the Declaration of Helsinki (2013).

#### Stimuli and task

Stimuli consisted of twelve unique Dutch words: *“waar”, “jip”, “ik”, “ga”, “boe”, “kom”, “hoed”, “spelen”, “plukken”, “allemaal”, “janneke”, “grootmoeder”*. These words were selected from a Dutch children’s book “Jip and Janneke” to maximize the occurrence of specific phonemes (/k/, /p/, /a/ and /u/) used in a different task^15^.

During the word production experiment, each participant was presented with a visual cue (target word on screen) and instructed to read aloud the word shown on screen. Each word was presented ten times. The order of words was randomized over participants, resulting in 120 word trials in total. Inter-trial interval duration was randomized and on average was equal to four seconds in S1, S2, S3 and S5: 4.08 ± 1.52 in S1, 3.93 ± 1.48 in S2, 4.11± 1.74 in S3, and 4.06 ± 1.34 in S5. Due to time constraints during awake surgery, inter-trial interval in S4 was reduced to 2.49 ± 0.36. Four of the participants implanted with ECoG grids for epilepsy monitoring (S1, S2, S3, S5) were also presented with “rest trials” (“-” on screen), during which they were instructed to be silent. The participant who underwent awake brain surgery (S4) was not presented with “rest trials” due to time constraints. The experiment took 9.18 minutes in S1, 9.09 minutes in S2, 8.96 minutes in S3 and 9.51 minutes in S5. S4 completed the task in 6 minutes and 2 seconds.

#### Experimental procedures

All participants were implanted with HD ECoG grids over the sensorimotor cortex. The suspected pathological regions did not extend to the sensorimotor region covered by the HD grids. This was clinically confirmed after implantation. In all participants, except for S2, HD grids were implanted over the left hemisphere. The hemisphere for HD ECoG implantation was the same as for clinical ECoG grids, and that was determined based on the clinical need. HD grid configurations differed slightly across participants with S1 implanted with 128 contacts, 1.2 mm exposed diameter, inter-electrode distance 4 mm, and S2, S3, S4 and S5 implanted with 128 contacts, 1 mm exposed diameter, inter-electrode distance 3 mm.

In the experiment with participants in the intensive epilepsy monitoring unit (S1, S2, S3 and S5), words were presented to the participants on a computer screen (21 in. in diagonal, at about 1 m distance) using Presentation software (Neurobehavioral Systems). In the experiment with the participant in the operating room (S4), words were presented on a tablet using custom Python scripts. In both cases, HD ECoG data were acquired using the NeuroPort neural recording system at a sampling rate of 2000 Hz (Blackrock Microsystems). Synchronization of the task with neural recordings was achieved based on event codes sent from stimulus presentation software to the recording computer.

In addition to the neural data, audio recordings were acquired using an external microphone (Audio-Technica AT875R), connected directly to the neural recording system. This ensured synchronization of the microphone recordings with neural data acquisition. Microphone data were recorded at 30000 Hz.

#### ECoG data processing

HD ECoG data were loaded in MATLAB using the NPMK toolbox. The data were preprocessed using custom software. First, all electrodes with noisy or flat signal (based on visual inspection) were excluded from further analyses (S1: none, S2: five electrodes, S3: 1 electrode, S4: 17 electrodes, S5: 1 electrode). Second, a notch filter was applied to the remaining electrodes to account for the effects of line noise (50 Hz and its harmonics). Third, common average (CAR) or bipolar referencing was applied. Fourth, data were transformed to the frequency domain using Gabor wavelet decomposition at either 60–300 Hz or 70–170 Hz in 1 Hz bins with decreasing window length (four wavelength full-width at half maximum). Fifth, high frequency band (HFB) amplitude was obtained by averaging amplitudes for the entire range of extracted frequencies. Sixth, the resulting time series per electrode were log-transformed and downsampled to 100 Hz. In total, four preprocessed HD ECoG files were made per participant: bipolar in the range of 60–300 Hz, bipolar in the range of 70–170 Hz, CAR in the range of 60–300 Hz and CAR in the range of 70–170 Hz.

In order to provide statistics of baseline HD ECoG activity, during the task, per participant, we identified a continuous fragment of at least five seconds in length, during which the participant remained silent and at rest. Mean and standard deviation of baseline neural activity was calculated per electrode. These values were used for normalization of HD ECoG task data during training of speech reconstruction models.

Electrode locations were coregistered to the anatomical MRI in native space using computer tomography scans^66, 67^ and FreeSurfer^68^.

#### Microphone data processing

Microphone data were loaded in MATLAB using the NPMK toolbox and resaved in the wav format at the original sampling rate of 30000 Hz. Event codes from the stimulus presentation software provided information about onset of each trial and the target word per trial, but to extract precise timing of the pronunciation of each word, we used the PRAAT toolbox (https://praat.org)^69^. Using PRAAT we manually annotated each participant’s microphone recording file (wav) with onsets and offsets of individual words.

Per participant, we also performed audio denoising procedures to obtain “clean” versions of each original, or “raw”, microphone recording. Denoising was performed with Audacity. First, a high-pass filter was applied at 60 Hz. Then, a built-in noise removal tool was used first to estimate background noise from a fragment of three to five seconds in length, and then to subtract the estimated noise signal from the rest of the audio. This process was repeated four or five times to obtain cleaner background. Finally, additional manual corrections were applied to remove irregular noise, such as occasional talking of staff present in the room, sounds of medical equipment or other background noise. Both raw and clean audio files were used in speech reconstruction.

For training of speech reconstruction models, we extracted spectrogram features from raw and clean audio files of microphone recordings. We extracted log-mel spectrograms in the frequency range of 80 to 7600 Hz, as they best approximate human speech perception. For parameter optimization, we varied the following spectrogram extraction parameters: number of mel frequency bins (40 or 80), window length (1470, 882, 525) and hop size (1470, 882, 294). The choice of the latter two parameters determined the sampling rate of the resulting audio: 15, 25 or 75 Hz, respectively.

### Speech reconstruction

#### Modeling approach

Brain data was used as input and speech spectrograms as output. Only data during speech pronunciation was used in model training and testing. Spectrograms of the microphone audio data the models and trained, tested and validated on are referred to as target audio spectrograms. The speech reconstruction model was trained to solve a regression problem of mapping 2d brain inputs (electrodes by time) onto 1d audio outputs (frequency vector at one time point):

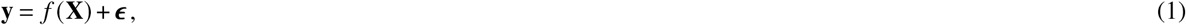

where **y** ∈ ℝ^*m*^ is a vector of spectrogram frequency values per one time point and *m* frequency bins, **X** ∈ ℝ^*n*×*p*^ is a matrix of associated neural recordings with *n* corresponding to the number of data points over time and *p* corresponding to the number of ECoG electrodes, *f*(·) is a non-linear transformation function and *ϵ* ∼ 𝒩 (**0**, *σ*^2^**I**) is a noise term. The transformation function is learned by the reconstruction model based on the artificial neural network. The reconstruction model is trained end-to-end.

#### Reconstruction model architectures

Three commonly used artificial neural network architectures were employed as the reconstruction models: a multi-layered perceptron (MLP), a densenet convolutional neural network (DN) and a sequence-to-sequence recurrent neural model (S2S, Figure 8). MLP is an artificial neural network that consists of layers of nodes that apply linear transformations to their inputs, followed by a non-linear activation function. In our configuration, each linear layer was followed by batch-normalization and leaky ReLU (negative slope= 0.25) activation function.

**Figure 8.**
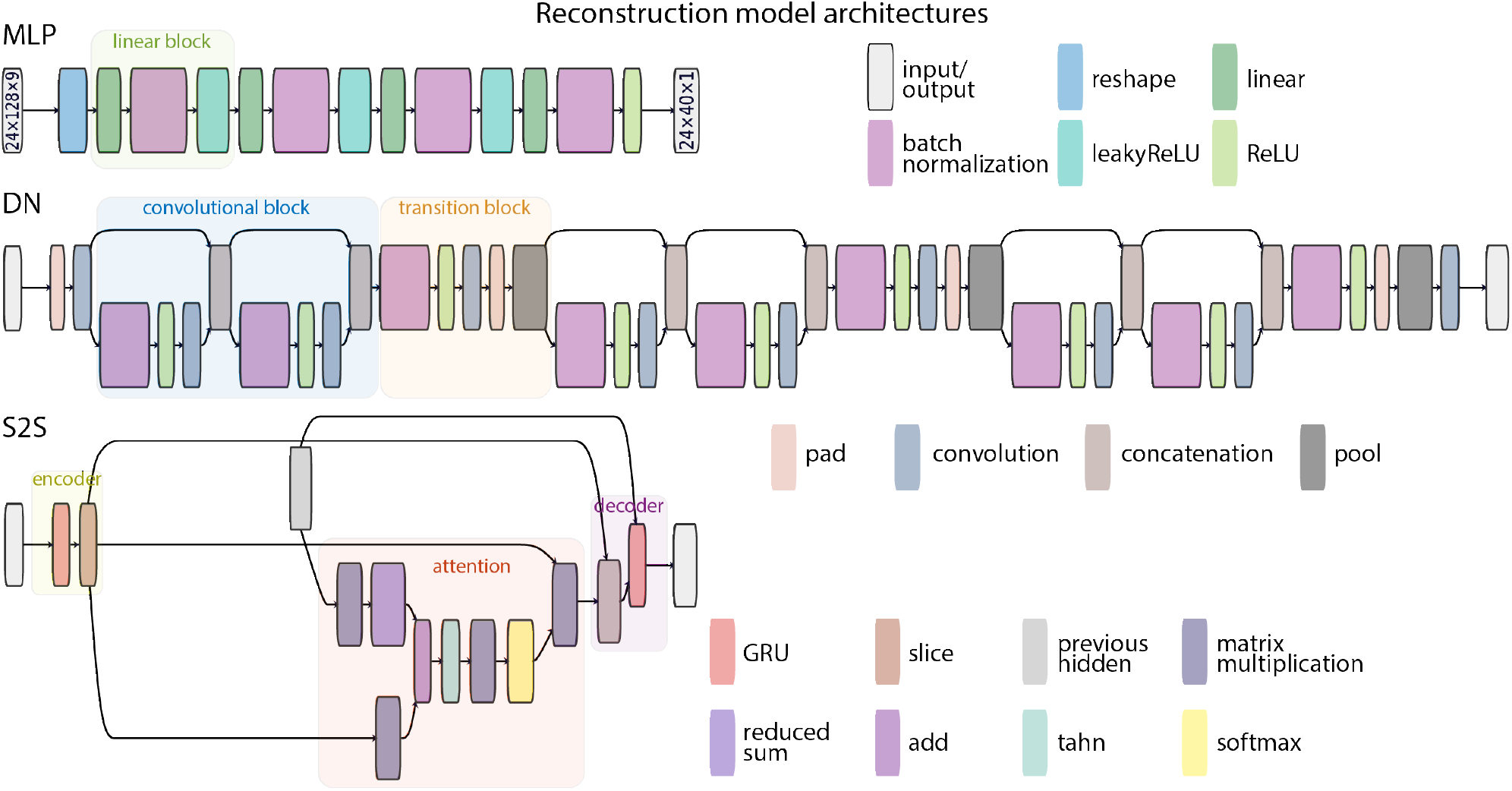
Reconstruction model architectures: MLP, DN and S2S. Visualization of model architectures was made with Netron (https://github.com/lutzroeder/netron) and subsequently modified. Specifically, for improved readability, layers with trivial operations, such as squeeze and transpose, were removed and the color scheme was extended to include unique colors for all individual layers.

DN^54^ is a convolutional neural network architecture with skip connections that aim to handle vanishing gradients. It consists of blocks of convolutional layers. In a block, the outputs of all preceding layers are directly connected as additional inputs to all subsequent layers via concatenation. To accommodate processing of growing inputs, each subsequent layer grows in number of nodes (channels) by a specified *growth factor*. To compress the model and downscale its complexity, transition blocks are inserted after each convolutional block. They reduce the amount of information currently processed by the network by a value of the *reduce* parameter. In addition, *bottleneck* layers can be used to reduce the number of network channels in a block (by using 1 × 1 convolutions).

S2S^55^ is a recurrent neural network architecture with an encoder and a decoder components. The encoder processes temporal information in the input, its representation in the hidden state at the last time point is used as input to the decoder. Additionally, an attention mechanism is often employed for better performance. Here, we used a form of global attention^70^ to weigh encoder outputs and produce a context vector that is used as input to the decoder. Both encoder and decoder components in our S2S are based on a gated recurrent units (GRU)^71, 72^.

#### Parameter optimization

Optimization was performed per model architecture and per subject using tree-structured Parzen estimators (TPE) as implemented in the Optuna (https://optuna.org) package^56^. This enabled us to optimize choices of categorical variables, yet came with a drawback of having to use independent samples. The latter meant that hyperparameter values were selected independently without considering interactions between them. The parameter optimization procedure aims to identify a set of model parameters *θ* that minimize the objective function *f* (*θ*) – reconstruction loss in our case:

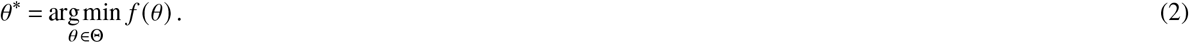

In order to identify optimal parameters, TPE estimate the posterior probability *p* (*y*| *θ*), where *y* is the validation reconstruction loss. This is done using the Bayes rule:

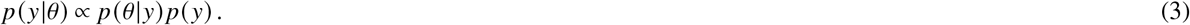

The probability *p* (*θ* |*y*) is estimated using a decision boundary based on a threshold *y*^*\**^:

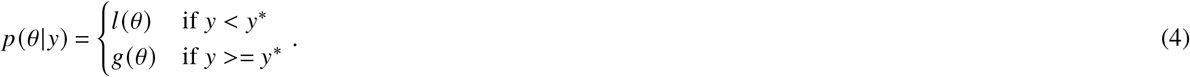

Effectively, TPE estimates two propability distributions of model parameters: *l* (*θ*) associated with *y* values below the threshold and *g*(*θ*) for *y* values above the threshold. The expected improvement *E I* of the optimization algorithm is proportional to the ratio *l* (*θ*) to *g* (*θ*). *E I* is maximized by iteratively drawing samples from *l* (*θ*) and *g* (*θ*), calculating their ratio and selecting a set of parameters *θ* that maximizes that ratio:

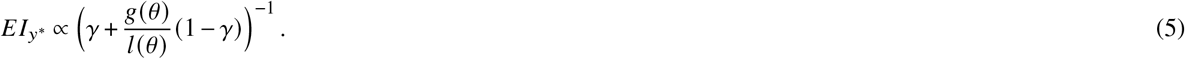

Using the TPE framework as implemented in Optuna, we optimized several input and output parameters and model hyperparameters. Separate input files were used for referencing (bipolar or CAR) and band width (60–300 or 70–170 Hz). The window of input data for reconstructing one time point of the audio could also be adjusted (160 or 360 ms). Separate output files could also be used depending on the number of mel features (40 or 80) and audio type (clean or raw). Depending on the target audio sampling rate (15, 25 or 75), corresponding audio and neural files were used.

Model parameters included architecture-specific and general parameters. General parameters included learning rate of the optimizer and dropout ratio. Parameters specific to the model architecture included number of layers and nodes per layer in MLP, presense of bottleneck layer, growth factor, reduce factor and number of layers in DN and number of encoding layers, number of decoding layers and bidirectionality in S2S.

#### Training procedures

Data was divided into train, test and validation sets in a pseudorandom fashion. One trial per each word was selected for validation (twelve words in total) and another trial for each word was selected for test (twelve words in total). Pseudorandomization ensured that no consecutive trials were used in validation or test. It also attempted to distribute validation and test trials such that they were well spread-out throughout the entire task.

Each model was trained to take in input brain data and reconstruct target audio spectrogram. To reconstruct one time point of the audio spectrogram, a temporal window of varying length (160 or 360 ms) around the target time point was selected in neural data as input to the model. Each model was trained for 500 epochs, after which the reconstruction loss was calculated on the validation set. Adam optimizer with fixed parameters except for the learning rate was used. All training and testing procedures were implemented in PyTorch (https://pytorch.org). The models were trained using a single graphics processing unit NVIDIA GeForce RTX 2080 Ti.

The described model training procedure was embedded in the model optimization study by Optuna. A separate study was made per model (MLP, DN, S2S) and subject (S1, S2, S3, S4, S5). Each Optuna study was set to run for 100 “trials” (not the same as the task trial, “trials” in an Optuna study refer to a single model training procedure with a fixed set of parameters). In each Optuna trial, a set of model parameters was selected, the model was trained for 500 epochs and the reconstruction loss on the validation set was calculated. The latter was used to guide parameter selection in the subsequent Optuna trials. In the end, the best Optuna trial with the corresponding parameter set was identified as the Optuna trial with the lowest reconstruction loss on the validation set.

#### Parameter importance

After model optimization, functional ANOVA (fANOVA)^73^ as implemented in Optuna, was used to fit a random forest regression model and predict the reconstruction loss in the validation set given a specific parameter set. FANOVA estimated the effects of main variables (parameters) and interactions between them on the variance of the dependent variable (reconstruction loss).

#### Testing procedures

To obtain reconstruction results on the test dataset, leave-one-out cross-validation (LOO-CV) was employed. A separate model was trained for 500 epochs per each word in the test set using the set of optimal parameters identified during model optimization. Data from the validation set was added to the training set to increase the amount of available data.

The model at the 500th epoch was used to obtain reconstructed spectrograms of the target test word. For all evaluation procedures, reconstructed and target spectrograms were computed per test word trial beginning with word onset and ending with word onset plus one second, which matched the duration of the longest trial across all subjects. An external vocoder, Parallel WaveGAN^74^ (as implemented here: https://github.com/kan-bayashi/ParallelWaveGAN), was used to synthesise speech waveforms from the reconstructed spectrograms. Parallel WaveGAN is a generative deep neural network trained to synthesize speech waveforms given a mel-spectrogram. We used publicly available weights of the vocoder model (ljspeech_parallel_wavegan.v1) pre-trained on the LJ Speech dataset of spoken English (LibriVox project). Our experiments showed that despite being trained on English speech the model synthesized high-quality intelligible Dutch speech as well. In order to synthesize waveforms both target and reconstruction spectrograms were upsampled to match input dimensions the vocoder model was trained on: 80 mel-frequency bins at a sampling rate of 86.13 Hz. Each evaluation metric (except for speaker classification and human perceptual judgments) was then computed per LOO-CV fold (per one of the 12 word trials in the test set).

### Low-level evaluation metrics

Pearson correlation, as implemented in the Scipy (https://scipy.org) library for Python, was computed between reconstructed and target spectrograms by vectorizing the spectrograms per word and correlating resulting vectors. This was done to account for the relative differences in spectrogram energy over frequency bins and time points.

Match in voice activity detection (VAD) and short-term objective intelligibility (STOI)^58^ were computed on waveforms synthesized from reconstructed and target spectrograms using the WaveGAN vocoder. VAD was applied separately to each audio file (reconstructed and target speech). It was computed per each 30-millisecond window of audio and resulted in a string of “one” (voice activity detected) and zero (no voice activity detected) values per file. We used python-based interface (https://github.com/wiseman/py-webrtcvad) for online free VAD service https://webrtc.org. VAD match was then calculated as a fraction of VAD output matches between reconstructed and target speech fragments:

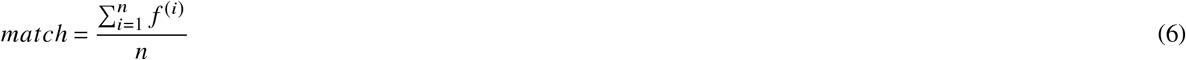

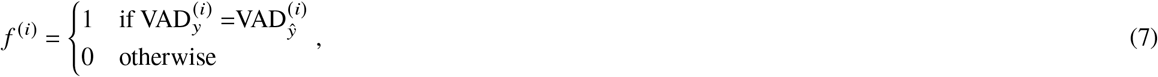

where *y* is the target waveform, *ŷ* is the reconstructed waveform of equal length and *n* is the number of 30-millisecond windows in each file, per which one VAD value was computed.

STOI was computed on synthesized waveforms using a STOI package for Python (https://github.com/mpariente/pystoi). For those waveforms with a duration that was shorter than the minimum required for STOI computation (30 frames, or 300 ms, recommended for intermediate intelligibility), the algorithm set the output value to 10^−5^. All such trials were excluded from subsequent analyses.

For each low-level evaluation metric, surrogate baseline distributions were computed for calculation of statistical p-values. This was done by shifting the neural data to time points corresponding to non-speech fragments 1000 times prior to obtaining test reconstructions and recalculating each low-level metric for each shift. This was done because one of the metrics, namely VAD match, only distinguished between speech and non-speech fragments and did not discriminate between individual words. Additionally, to provide another baseline for Pearson correlation and STOI metrics that focused on distinctions between individual words, we computed statistical p-values based on word permutations. For this, we used 1000 permutations of word labels when selecting data from the target audio files, thereby disrupting the correspondence between reconstructed and target speech fragments. The statistical p-values were comparable to those obtained with the shift-based baseline.

### High-level evaluation metrics

#### Machine learning classifiers

Word classification was performed using logistic regression. Per each test word (out of 12, in the LOO-CV), the classifier was trained either on raw brain input (same data as input to the speech reconstruction models), or target audio spectrograms from the train set data. The classifiers were then tested on the test data: raw brain data or reconstructed spectrograms (obtained by passing raw brain input through the speech reconstruction models).

In addition to this, a separate noise study was performed to estimate the performance robustness of word classifiers. We iteratively added incremental amounts of white noise to the classifier input data during training and recomputed their performance on test data. In the case of the reconstructed data, the noise was applied after the reconstructions were obtained with the deep learning models.

Next, a speaker classifier was trained on target speech spectrograms of all participants. Per speech reconstruction model, it was tested on the reconstructed spectrograms. Python library Scikit-learn (https://scikit-learn.org) was used for training and testing all classifiers.

For both metrics, permutation tests (1000 shuffles of classification labels) were conducted to provide a surrogate baseline distribution for calculation of statistical p-values for each observed metric.

#### Human perceptual judgments

Human perceptual judgments of reconstructed spectrogram were collected from healthy volunteers in a series of online behavioral experiments. We conducted three experiments: a word recognition experiment (I), a speaker recognition experiment (II) and an audio comparison experiment (III). Participants (native Dutch with normal hearing; I: 30 participants, age 27 ± 8, 12 females; II: 29 participants, age 27 ± 8, 7 females; III: 29 participants, age 27 ± 8, 8 females) were recruited online via Prolific (https://prolific.co). Each experiment was listed separately, and Prolific users were free to choose to participate in any number of experiments in any order. All participants gave their written informed consent and were reimbursed for their participation. The study was approved by the ethical committee of the Faculty of Social Sciences at Radboud University.

Experiments were implemented using Gorilla Experiment Builder (https://gorilla.sc). Because target and reconstructed microphone recordings were used in the experiments, to make voices of individual participants (S1, S2, S3, S4 and S5) unrecognizable, pitch of all audio files was shifted by 1 or 2 tones up or down, depending on the ECoG participant. In each experiment, behavioral study participants were told that the study investigated the effect of audio degradation on speech intelligibility. In experiment I, participants were presented with an audio fragment and two written words, and were instructed to select the word that corresponded to the audio they heard. In experiment II, participants were presented with three audio fragments: the “target” audio at the top of the screen, and two audio fragments at the bottom, labeled “speaker X” and “speaker Y”. Participants were instructed to select the speaker label that corresponded to the speaker of the “target” audio. In experiment III, participants were presented with two audio fragments and were instructed to select the fragment with an overall better audio quality (more intelligible, less noisy). Data from each experiment, as well as screenshots of each experiment’s layout have been made publicly available (see Code and data availability). In experiments I and II, synthesized audio waveforms from both reconstructed (MLP, DN and S2S) and target spectrograms were used. Only optimized models were used for producing reconstructions, resulting in 360 trials: data from 5 ECoG subjects × 12 words from the test set × 3 speech reconstruction models × 2 types of audio files (reconstructions and targets). In experiment III only reconstructed audio were used for direct comparison of perceptual quality of MLP, DN and S2S outputs. In addition, both optimized and non-optimized (that used a randomly chosen parameter set from the first Optuna “trial”) models were used for producing and comparing the reconstructions, resulting in 360 trials: data from 5 ECoG subjects × 12 words from the test set × 3 speech reconstruction models × 2 types of optimization (optimized and non-optimized).

Behavioral judgments from experiments I and II were loaded into Python, response accuracy was averaged over test words, and the results were plotted per behavioral participant, HD ECoG subject and model (Figure 6a, b). Responses from experiment III were assigned an arbitrary weight of 1 if reconstruction with the optimized model was rated as having higher quality, and a weight of -1 otherwise. The weights were then averaged over test word trials and plotted per behavioral participant, HD ECoG subject and model. Non-parametric tests were used for calculation of significance. For this, on every trial, instead of taking the participant’s response, the choice was made at random. This procedure was repeated 1000 times. The surrogate distributions were used in calculation of statistical p-values for the observed human judgments.

Finally, we made scatter plots and fitted a regression line between results obtained with low-level evaluation metrics (Pearson correlation, VAD match and STOI) and human perceptual judgments.

### Input perturbation analysis

To estimate individual electrode contributions to the performance of the reconstruction model, an input perturbation analysis was performed. For this, per subject and model, we used the reconstruction loss of the optimized trained model on test data as our reconstruction loss baseline. Then, we iteratively recomputed the reconstruction loss on test data the number of times equal to the number of electrodes. Each time, prior to computing the loss but after input normalization, we zeroed out input of one of the electrodes. Then, we calculated the difference between the resulting reconstruction loss compared to the baseline loss value (when input data from all electrodes was used). This way, if an electrode contributed to the reconstruction, taking out its signal during testing resulted in increase of the loss compared to the baseline. We z-scored the resulting difference values and plotted them on the brain.

### Audio contamination analysis

Recent work has demonstrated that direct neural recordings can show signs of acoustic contamination in their acquired signals^75^. To account for this possibility we performed an audio contamination analysis. Similar to the study by Roussel et al., we computed correlations between spectrotemporal signals of microphone speech recordings and HD ECoG activity. Spectrograms were computed using the librosa (https://librosa.github.io/librosa) package. For this, short-time Fourier transform (STFT) was applied to audio (at sampling frequency of 16kHz) and brain (at sampling frequency of 2kHz) data. We selected signal processing parameters that produced spectrotemporal signals of matching temporal resolution (STFT window of 131 ms for audio and STFT window of 256 ms for HD ECoG), necessary for the correlation analysis. A default window overlap of 25% and a Hann window function were used. Prior to STFT calculation, HD ECoG data were notch-filtered to account for the line noise (at 50 Hz and all its harmonics) and referenced using either the common average or the bipolar scheme, since both types of signals had been used in the previous analyses. Pearson correlation coefficient was computed for audio and HD ECoG signals across all speech production trials (between 116 and 120 trials depending on the participant).

The analysis did not reveal any signs of acoustic contamination in four subjects out of five (Figure 9). Only S5 showed noticeable correlations between several HD ECoG channels and the audio spectrogram in the range of 100–140 and 210–280 Hz. To perform a more detailed analysis we used the acoustic contamination toolbox provided by Roussel and colleagues^75^. Using the toolbox, we calculated cross-correlations between audio and HD ECoG signals of S5 during speech and calculated the statistical p-value for audio-ECoG correlations along the main diagonal. The detailed analysis revealed that 1) the bulk of the correlation peaked at the negative lag of 170 milliseconds (ECoG activity preceded audio), and 2) the mean diagonal correlation of .2 did not reach significance based on the permutation tests (with the statistical p-value of *p* = .19. based on 10000 reshuffles of contamination matrices).

**Figure 9.**
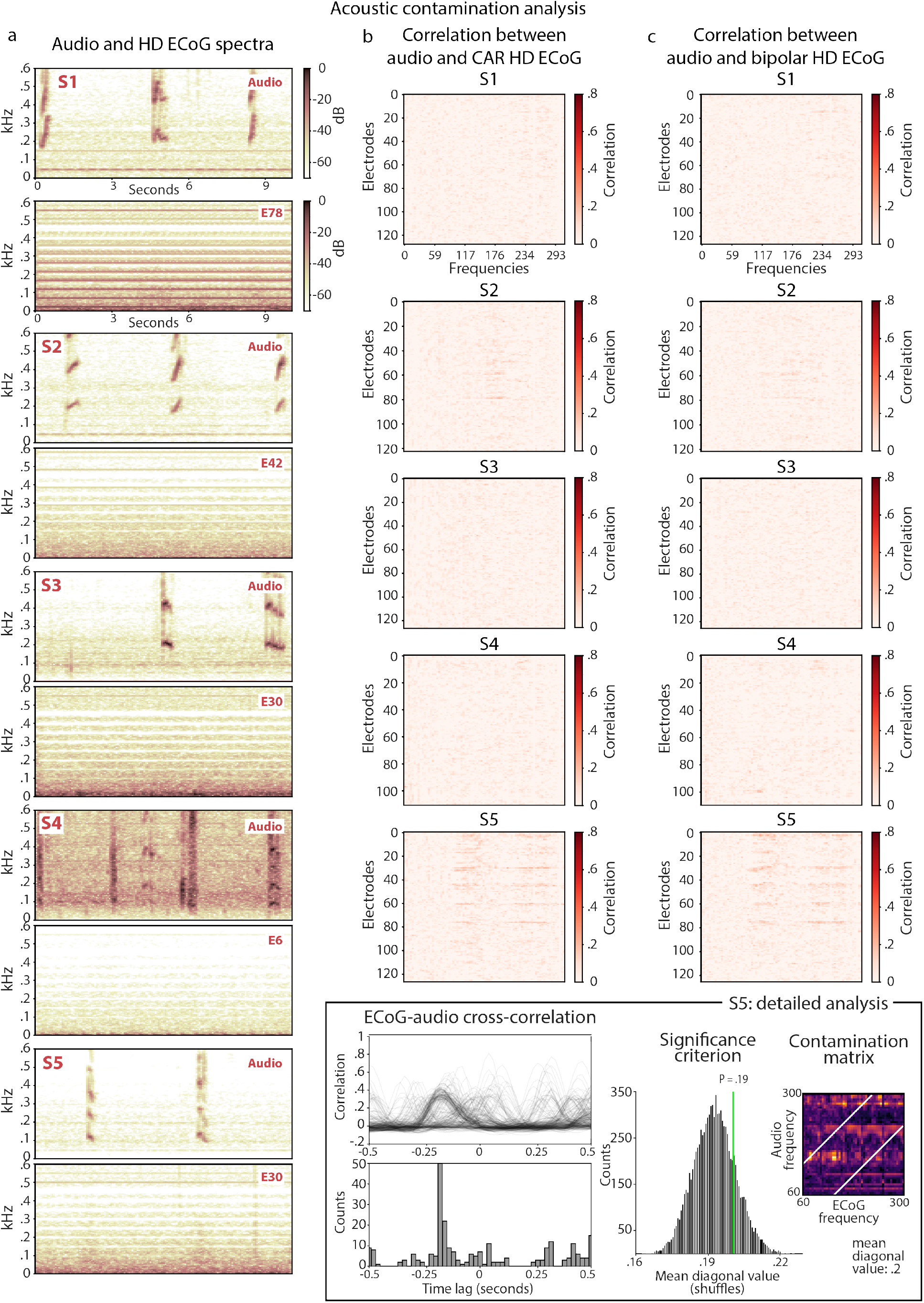
Results of the acoustic contamination analysis. a. Spectrograms of the microphone audio recordings and associated HD ECoG signals. Per subject ECoG signal of one electrode is shown. The electrode was selected based on t-tests that compared mean HFB activity between speech and silence trials per electrode. The electrode with the largest positive t-value was used for the plot. Selected parameters resulted in the matched temporal resolution of the spectrogram signals (16000 / (4096/ 4) in audio and 2000 / (512 / 4) in HD ECoG). The audio spectrogram is shown in the frequency range that matches the ECoG spectrogram. Plots show HD ECoG data prior to referencing (common average or bipolar). b-c. Correlation plots between audio and HD ECoG temporal signal per frequency bin (between 0 and 300 Hz) and per electrode. Each value is the average Pearson correlation over all word production trials. Correlation plots are shown for CAR (b) and bipolar (c) reference schemes for HD ECoG signals. The bottom right panel shows results of the detailed contamination analysis using a toolbox from Roussel and colleagues^75^. The panel shows results of the cross-correlation analyses that indicate that the bulk of the correlation was not centered around zero (when ECoG and audio signals are aligned in time), but was shifted towards the lag of -170 milliseconds. Negative shifts mean that ECoG activity preceded audio data it was correlated with, which agrees with the previous reports of pre-activation of motor and premotor cortices prior to speaking. The formal analysis revealed that mean correlation between audio and ECoG data was .2, which did not reach significance based on the permutation tests (with the statistical p-value of *p* = .19. based on 10000 reshuffles of the correlation matrices).

### Code and data availability

Code of all analyses can be accessed at https://github.com/Immiora/word_decoding_HD_ECoG. Data from human perceptual judgments can be found at https://doi.org/10.34973/c07k-v019. HD ECoG data will be released pending a temporary embargo and required de-identification procedures.

## Acknowledgements

This work was supported by the European Research Council (Advanced iConnect Project Grant ADV 320708) and the Netherlands Organisation for Scientific Research (Language in Interaction Consortium, Gravitation Grant No. 024.001.006; and INTENSE Consortium, Grant No. 1761). We thank Frans Leijten, Cyrille Ferrier, Geert-Jan Huiskamp, Sandra van der Salm and Tineke Gebbink for help with collecting ECoG data; Peter Gosselaar and Peter van Rijen for implanting the electrodes; ECoG patients and participants of the behavioral experiments for their time and effort; Zuzanna Dedyk for help with implementing behavioral experiments; and the members of the UMC Utrecht BCI team for help with ECoG data collection.

## Author contributions statement

All authors conceived the idea of the study and the approach and contributed to data acquisition, J.B. preprocessed the data, implemented the modeling approach, performed the analyses and wrote the initial draft of the paper. All authors reviewed the manuscript.

